# Standardized human tests show that most vision models are face-blind

**DOI:** 10.64898/2026.09.18.750666

**Authors:** Kushal Dudipala, Florencia Martinez-Addiego, Dobromir Rahnev, Bradley C. Duchaine, N. Apurva Ratan Murty

## Abstract

Face recognition ability varies enormously across humans. Individuals with face-blindness (prosopagnosia) struggle to recognize even close family members while super-recognizers can identify strangers with exceptional accuracy. Where do ANNs fall within the human face recognition spectrum? Here, we administered the standardized tests used to characterize human face recognition ability to a diverse set of models, allowing us to contextualize a model’s performance within the distribution of human behavior. We found that the majority (55%) of models to be classified as face-blind and that even the best face-trained models do not cross the human threshold to be considered a super-recognizer. Interrogating the internal representations of these models showed that models that performed well on standardized face recognition tasks were more identity-selective and viewpoint-invariant. We also found that low-performing models did contain some identity information in independent representational subspaces. Removing viewpoint-dependent subspace improved face recognition abilities in 49 of the 53 models tested. Targeted unit ablations further identified opposing contributions, with viewpoint-dependent units disrupting identity coding and viewpoint-tolerant units supporting it. Together, our results show that most AI models are face-blind with worse face recognition ability than humans, and demonstrate how differences across AI models can be used generate testable hypotheses about the computational basis of human face recognition in humans which can then be probed in future studies.

## 1 Introduction

Humans vary widely in their ability to recognize faces. At one extreme, super-recognizers can identify even unfamiliar people with remarkable accuracy. At the other extreme, individuals with prosopagnosia (face-blindness) struggle to recognize even familiar people from their faces. In between lies the broad range of face recognition abilities of the ‘neurotypical’ population. Decades of behavioral research has developed standardized behavioral paradigms for measuring these differences in face recognition abilities in humans (Benton et al. [1983]; Warrington [1984] Duchaine et al. [2007]; Duchaine and Nakayama [2005]; Duchaine and Nakayama [2006]).

Where do contemporary AI models, including our leading face recognition systems, fall within this landscape? With the rise of AI, artificial neural networks (ANNs) have also become very good at recognizing faces, with some reports of these systems now exceeding average human performance on standard benchmarks (Schroff et al. [2015]; Sun et al. [2015]; Taigman et al. [2014]). But these benchmarks were developed to evaluate machine performance and comparisons to the ‘average’ human, and consequently mask the actual variability seen in humans ([Fung et al., 2026]). Moreover, to our knowledge, contemporary ANNs have not been systematically evaluated on standardized human face recognition tests (e.g., the Cambridge Face Memory Test; CFMT; Duchaine and Nakayama [2006]). The CFMT is used to measure and classify face recognition ability in humans. Do ANN models evaluated on the CFMT behave like neurotypical observers, approach the extraordinary abilities of super-recognizers, or show the difficulties associated with prosopagnosia? Beyond benchmarking, placing models along the human spectrum could provide experimentally accessible systems for studying the mechanisms underlying individual differences in human face recognition abilities (Fung et al. [2026]). Models that resemble super-recognizers or individuals with prosopagnosia can then be directly interrogated to find the computational and representational differences associated with exceptional or impaired recognition, giving researchers new testable hypotheses about the neural mechanisms underlying such differences in humans (Schrimpf et al. [2020]).

Our key contributions are threefold. *First*, we place contemporary vision models directly on the human spectrum of face recognition ability using standardized behavioral tests. *Second*, we identify a representational signature that predicts face recognition ability: high-performing models preserve face identity information across changes in viewpoint, whereas poorer models are more strongly organized by viewpoint. *Third*, through targeted subspace projections and unit ablations, we show that identity information can be present but obscured by viewpoint-dependent structure, and identify components that make opposing contributions to face-identity coding. Together, these results show how interrogating models with distinct behavioral phenotypes can generate testable hypotheses about the computational basis of individual differences in human face recognition Fung et al. [2026].

## 2 Methods

We evaluated 53 models spanning diverse architectures, training regimes, and objectives (Supplementary Table 1) on three versions of the Cambridge Facial Memory Test (CFMT): CFMT (Duchaine and Nakayama [2006]), CFMT+ ([Russell et al., 2009]), and CFMT-AUS ([McKone et al., 2011]). Models were evaluated without training or fine-tuning by extracting their final embeddings prior to the classifier head. For vision-language models (VLMs), we used embeddings from the image tower. The CFMT variants are diagnostic tests of human face recognition ability, designed to measure differences between face-blind individuals, neurotypical observers, and super-recognizers. The experiments used the original test stimuli and closely followed the human procedure, consisting of an instruction phase, a learning phase, and a test phase. During learning, participants study target identities across three viewpoints; during testing, they identify the target from three candidate faces (chance accuracy = 1/3).

**Table 1:**
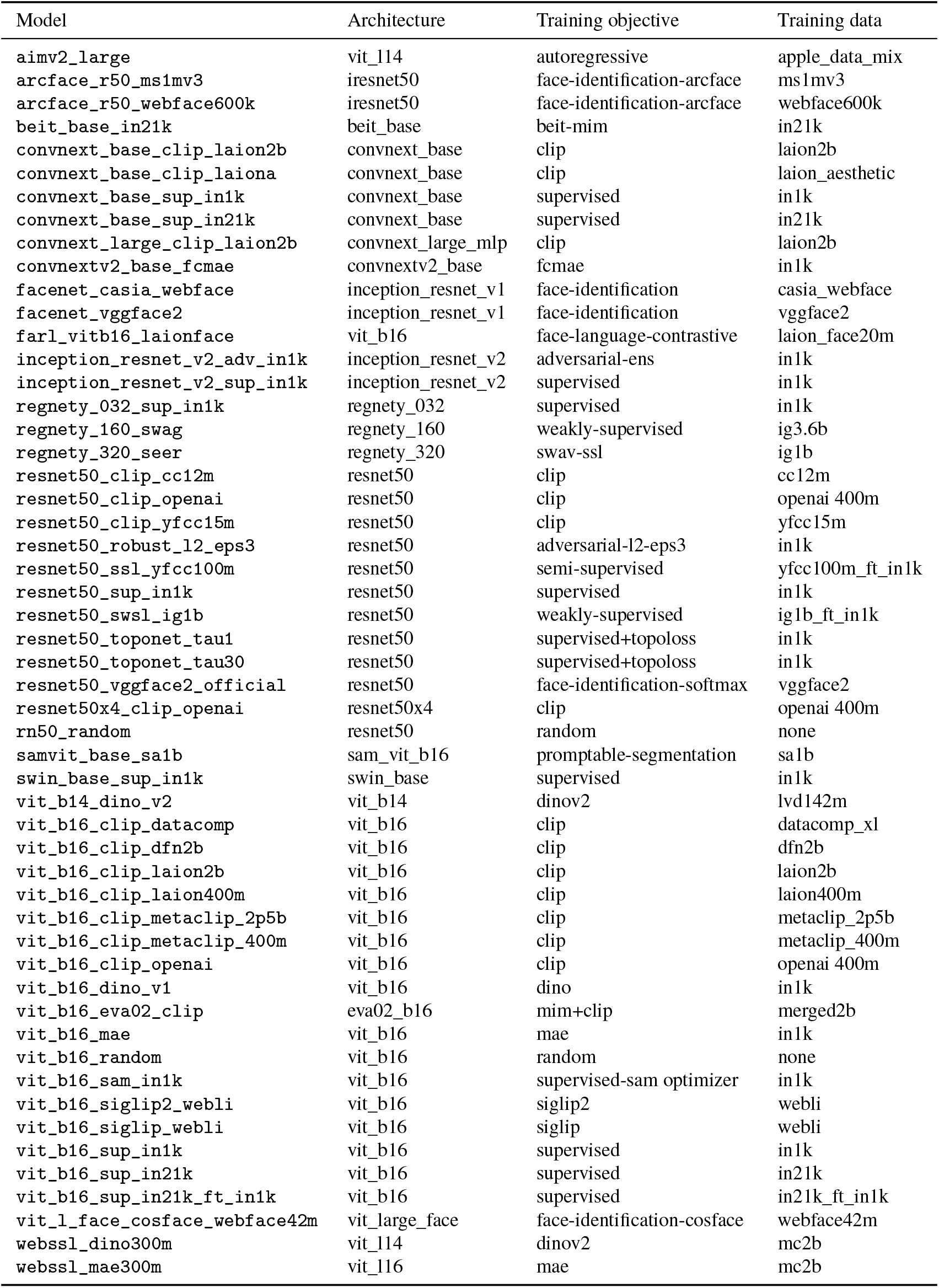
The 53 image encoders evaluated in this study.

**Table 2:**
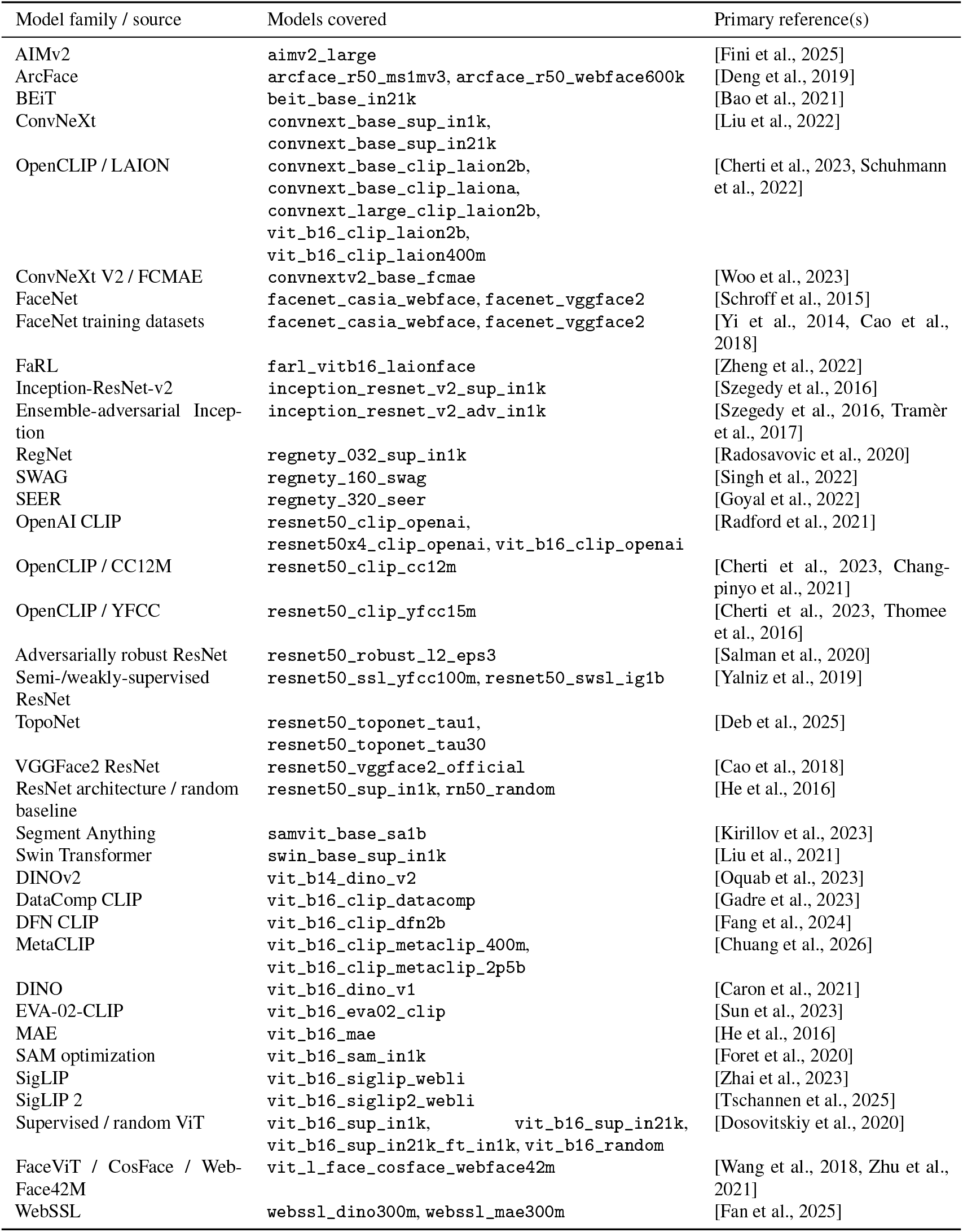
Primary references for the image encoders evaluated in this study.

Because models lack episodic memory, we replaced the learning phase with a face bank containing reference images for each target identity (18 images total: 6 identities × 3 viewpoints). For each test trial, candidate face embeddings were compared with reference embeddings using mean cosine similarity, and the candidate with the highest similarity was selected as the model prediction. Test scores were computed using held-out images to avoid performance driven by image-level matching.

Successful CFMT performance requires both viewpoint invariance and identity selectivity. We calculated viewpoint invariance and identity selectivity scores for each model (see Technical Methods). We then projected the subspaces associated with viewpoint dependence and identity selectivity out of the model’s representation (cross-validated; balanced; see Technical Methods). Finally, we identified strongly-associated viewpoint-dependent and viewpoint invariant units and progressively ablated them while examining the effect of their ablation on face recognition score (see Technical Methods).

## 3 Results

### 3.1 Standardized tests reveal that most AI models are face-blind and none are super-recognizers

We first sought to place AI models along the spectrum of human face recognition ability. We tested 53 models (see Methods) on three widely used human face recognition tests, the CFMT, CFMT-AUS and CFMT+. The CFMT and CFMT-AUS probed whether model performance fell within the range associated with prosopagnosia (face-blindness) or instead within the range of neurotypical observers. The more challenging CFMT+ instead tested whether models reached the range associated with super-recognizers (see Methods). We evaluated models on the same stimuli used in humans. Whereas humans provided explicit identity judgments, model responses were derived from the cosine similarity between representations at the final behavioral readout stage. Model responses were scored identically to human responses, allowing model performance to be directly compared with established human distributions and cutoffs.

On the classic CFMT task, model performance varied widely with a mean accuracy of 63% (**Figure 1a**). Published human data report mean human performance of 81% on this task [Stehr et al., 2025]. Individual-level neurotypical scores were not available for CFMT, precluding a direct statistical comparisons (though see next). Nonetheless, 29/53 models (55%) were below the 61% cutoff used to diagnose prosopagnosia in humans (Stehr et al. [2025]), which is much higher than 2–2.5% typically *×* observed in the population ([Bowles et al., 2009]). We observed a similar pattern with an independent set of faces in the CFMT-AUS. Model performance again varied widely, but the mean model accuracy was only 59% (**Figure 1b**) compared to 79% in humans McKone et al. [2011] (Mann-Whitney U, *p* = 4 × 10^*−*11^.) Are these differences in model performance consistent across tests, or do they instead capture distinct aspects of face recognition behavior? Model scores were strongly correlated *×* across the CFMT and CFMT-AUS (Spearman *ρ* = 0.8, *p* = 2 × 10^*−*15^), suggesting performance was robust across face recognition tests. Performance across these two tests was substantially lower than the corresponding human performance, with a much larger proportion of models falling within the range associated with prosopagnosia in humans (Bowles et al. [2009]).

**Figure 1:**
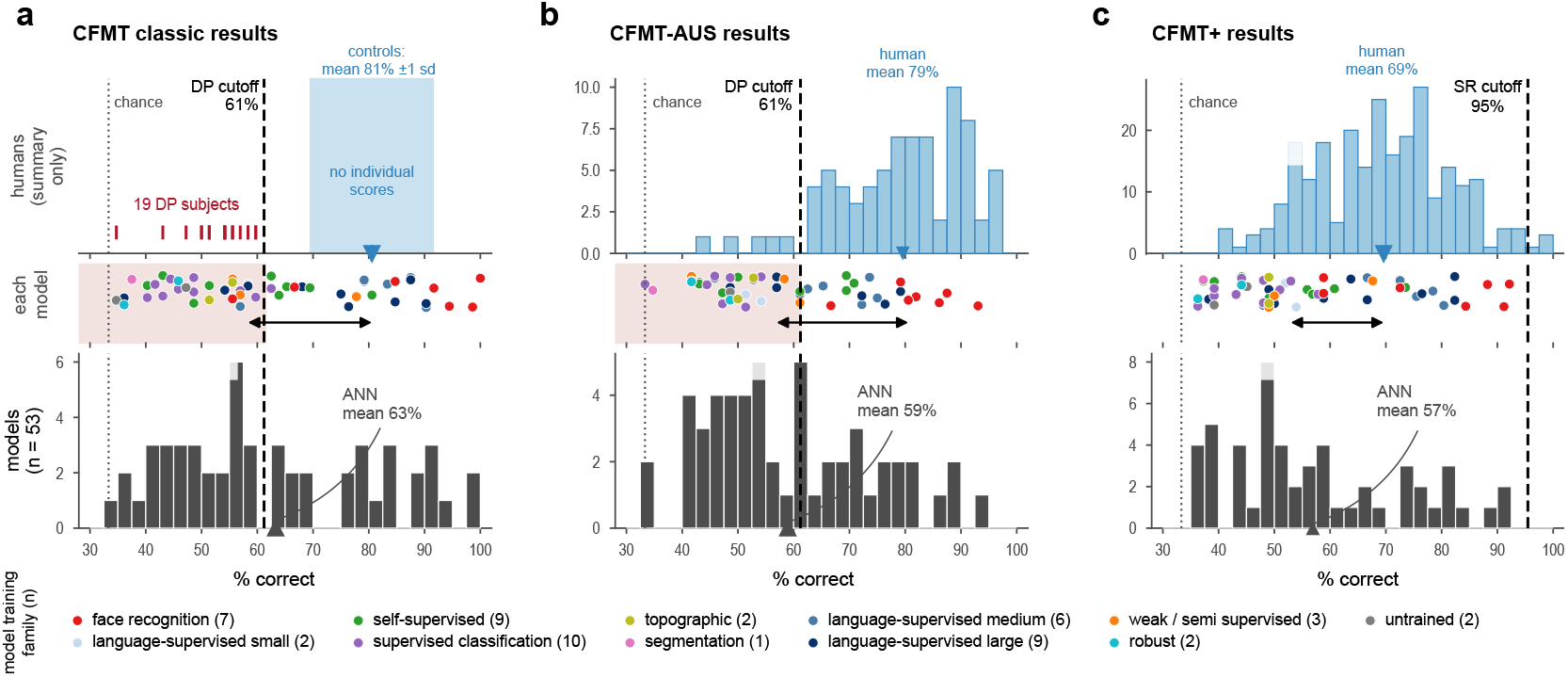
Face recognition performance of 53 models across three CFMT variants. **(a–c)** Corresponding results, with human data (top), individual model scores colored by training family (middle), and the model distribution (bottom) on a shared percent-correct axis. Blue marks human means; red ticks in (a) show 19 DP subjects. Dashed lines give the prosopagnosia cutoff (61%; a, b) and super-recognizer cutoff (95%; c). Model means fall near or below the DP cutoff across tests, and none reach the super-recognizer threshold.

Finally, we asked whether the best models reached the opposite extreme of the face recognition ability spectrum associated with super-recognizers. Here, we tested models on CFMT+. As before, model performance spanned a wide range (model mean = 57% (**Figure 1c**); human mean = 69% Bobak et al. [2016], Mann-Whitney U, *p* = 1.3 × 10^*−*7^). But critical to our test, not a single model reached the 95% cutoff used to classify super-recognizers in humans (0/53 models). Thus, while our models extended substantially into the range associated with impaired face recognition, even the best AI models trained explicitly on human face discrimination (face-trained models, red dots) do not have the exceptional levels of face recognition abilities observed in human super-recognizers.

Across two independent tests, the majority of models were classified within the range associated with humans with face-blindness, while the remaining models occupied the lower neurotypical range. Further, not a single model (including those trained for face recognition) reached the cut-off associated with super-recognition in humans.

### 3.2 Identity and viewpoint compete to structure face representations for recognition

What makes some models better at face recognition than others? Successful face recognition requires solving two related problems (1) distinguishing between identities (identity selectivity) and (2) having invariance across views (viewpoint invariance) [Murty and Arun, 2017, Ratan Murty and Arun, 2018]. We therefore asked how identity and viewpoint were organized within each model’s representation and quantified these properties from each model’s representations (see Methods). In high-performing face-recognition models (e.g., Facenet), different viewpoints of the same identity clustered together producing representation dominated by identity (**Figure 2a**, top). In a worse model (webssl-mae), the same data were spread by viewpoint to produce a view-dependent representation (**Figure 2b**, top). To quantify these effects we asked the extent to which these representational differences could account for variation in face recognition abilities across models. Individually, viewpoint invariance had a clear effect on a model’s CFMT score (Spearman *ρ* = 0.63, *p* = 5 × 10^*−*7^), while identity selectivity had a weaker affect on CFMT score (Spearman *ρ* = 0.24, *p* = 8 × 10^*−*2^). In addition, the terms had a covariance of 0.86, implying they capture overlapping variation on how models represent faces. To account for this overlap, we fit a regression model predicting each model’s CFMT score. These two properties explained 64% of the variance in CFMT performance across the 53 models (*R*^2^ = 0.64, *p* = 8 10^*−*12^, **Figure 2c**). Thus, differences in CFMT performance could be explained by differences in internal representational geometry.

**Figure 2:**
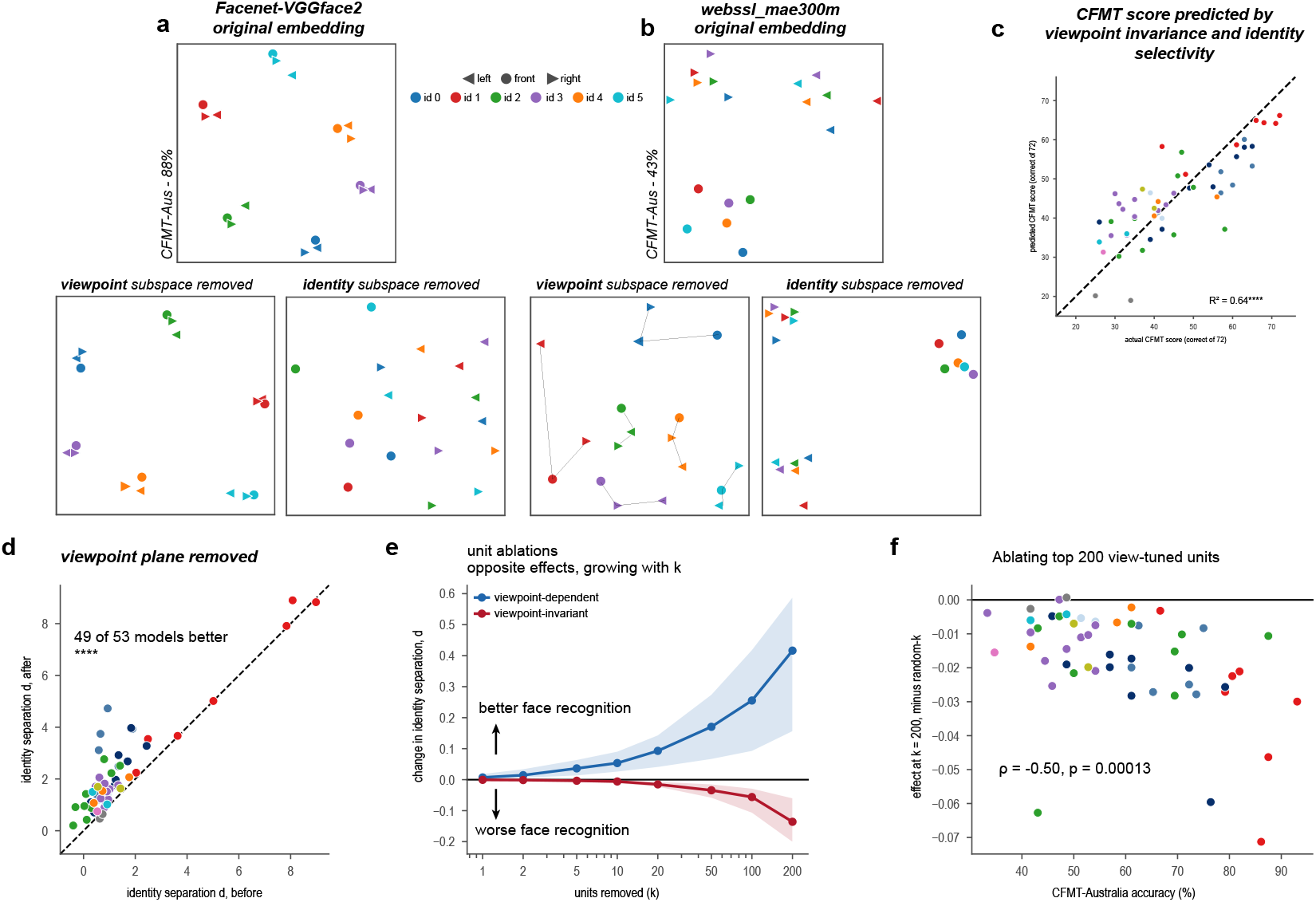
Identity and viewpoint compete to structure face representations. **(a,b)** MDS of the 18 face-bank embeddings for a good (a, top; Facenet-VGGface2 [Cao et al., 2018], 88%) and a weak model (b, top; webssl_mae300m [Fan et al., 2025], 43%) at the CFMT-AUS task. Bottom, MDS after removing the viewpoint or identity subspace. **(c)** CFMT score predicted from viewpoint invariance and identity selectivity (*R*^2^ = 0.64). **(d)** Identity separation *d* before vs. after viewpoint-subspace removal; 49/53 models lie above the diagonal. **(e)** Change in identity separation as units are ablated (*k* = 1–200), removing viewpoint-dependent (blue) or viewpoint-invariant (red) units. **(f)** Effect of ablating the top view-tuned units (*k* = 200, relative to a random-*k* baseline) against CFMT-AUS accuracy (*ρ* = *−*0.50, *p* = 1.3 *×* 10^*−*4^).

### Removing viewpoint structure exposes the latent identity structure in low-performing models

So far, these results do not explain why strong viewpoint structure accompanies poor identity coding. One possibility is that low performing models simply do not have any identity information. Alternatively, identity information may be present, but obscured by viewpoint-dependent representations. If the latter is true, then removing viewpoint-related variation should reveal a stronger identity structure. We tested this idea by finding the subspace associated with viewpoint and projecting it out of each model’s representation in a balanced cross-validated manner (estimating the subspace on 3 identities and evaluating on held-out identities). The same examples in **Figure 2a,b** (bottom) show this effect. While removing the viewpoint structure had relatively little effect on the identity-dominated Facenet model, this same manipulation significantly reorganized the webssl-mae model. Removing the viewpoint subspace improved the identity separation (better face identity recognition performance) in 49/53 models (**Figure 2d**). In contrast, removing the identity subspace made the representations viewpoint specific. This analysis shows that while some face identity recognition ability was present in these models it was obscured by competing viewpoint dependent representations.

### Targeted ablations reveal opposite contributions to face identity code

Can this competition be traced back to a model’s individual units? We identified units that were both strongly viewpoint-dependent and strongly viewpoint invariant and removed increasing numbers of each type. The two manipulations produced opposite effects: while removing viewpoint-dependent units progressively improved face identity abilities, removing view-invariant units progressively lowered face recognition abilities. These results move beyond identifying a causal role for these units by quantifying the functional consequences of their selective ablation within the broader neural population. Finally, we asked whether the reliance on these units was related to the model’s face recognition ability. Ablating viewpoint invariant units caused larger impairments in models that performed better on CFMT *− ×* (*ρ* = 0.5; *p* = 1.3×10^*−*4^) (**Figure 2f**). Thus, better face recognition models appear to depend more strongly on units that preserve identity across viewpoint changes. Together, these results provide a mechanistic account of the representational differences associated with face recognition ability in these models. Most AI models have poor face recognition abilities because their representations are dominated by viewpoint variation, though they contain some latent face identity structure. Better models rely on view-invariant components that preserve identity changes across pose.

## 4 Discussion

Where do ANN models fall on the spectrum of human face recognition ability? Despite their strong performance on conventional benchmarks, we found that, in stark contrast to the incidence of prosopagnosia in the human population (Bowles et al. [2009]), most models performed near or within the range associated with prosopagnosia. Moreover, no model reached the human super-recognizer range. This suggests that standardized human tests expose limitations in current AI face recognition models. Interrogating the models further revealed a potential computational basis for these differences. Better face recognition models organized representations primarily by face identity and preserved identity across changes in viewpoint (viewpoint invariant), whereas poorer models were dominated by viewpoint information (viewpoint dependent). We also found that even low performing models do contain some identity information organized in distinct representational subspaces. Removing the viewpoint-related subspace increased face identity separation in 49 of 53 models. Targeted ablations of individual units further showed opposing contributions of these components: removing viewpoint-dependent units improved face-identity recognition, whereas removing viewpoint-invariance units impaired it. That is, better models depended more strongly on viewpoint-invariance and were dominated by such units.

Together, these findings suggest that successful face recognition depends on separating identity from the transformations under which it must remain stable (in case of CFMT, viewpoint). Results from models may generate new predictions for humans: poor face recognition, including prosopagnosia, may reflect identity representations that are present but insufficiently disentangled from viewpoint. More broadly, our results illustrate how models with different behavioral phenotypes can be inter-rogated to generate testable hypotheses about the computational basis of individual differences in human cognition (Schrimpf et al. [2020]; Fung et al. [2026]).

## Technical appendices and supplementary material

### Model evaluation

Each three-alternative test trial is spliced into its candidate faces *x*_1_, *x*_2_, *x*_3_, one of which (index *k*) is the studied target. The face-bank reference images, *r*_1_, *r*_2_, are the target identity views, excluding the current trial view. This prevents low-level pixel correlations between the same faces from biasing the results. We pass each candidate face through the frozen encoder *ϕ*, along with the face-bank reference images for the target’s identity, and score each candidate by the mean cosine similarity between its embedding and those of the references:

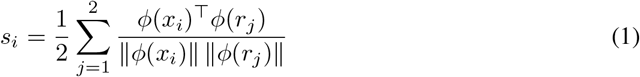

The face with the highest score is stored as the model’s “decision” for the trial. The trial is considered correct if this is candidate *k*. Since the model has three test options, chance is considered to be 1/3.

### Face memory benchmark

Models were evaluated on three variants of the Cambridge Face Memory Test (CFMT): the classic 72-trial version, the 72-trial Australian version, and the 102-trial long form. Because the variants differ in trial count, all scores are expressed as percent correct. Every trial is a three-alternative forced choice, so chance is 33.3% throughout. Human performance was taken from published samples: *n* = 78 subjects for the CFMT-Australia [McKone et al., 2011] and *n* = 253 Bobak et al. [2016] for the long form, both unselected, and for the classic variant 19 individually reported developmental prosopagnosic subjects with controls summarized only as a mean and standard deviation [Stehr et al., 2025].

### Linear viewpoint subspace

To quantify viewpoint invariance and identity selectivity for CMFT scores, we probe each model’s embedding space directly. From the CFMT-classic face-bank images we take embeddings *e*_*i,v*_ = *ϕ*(*x*_*i,v*_) of each face-bank image *x*_*i,v*_, for all 6 identities *i* and 3 viewpoints *v*. From this we form two families of pairs: same identity across views (_view_) and different identities within a view (_id_). We pool all pairwise cosine distances *d*(*a, b*) into one distribution with mean *µ* and standard deviation *σ*, and return two indexes as the mean *z*-scored distance for each family: Identity Selectivity = mean

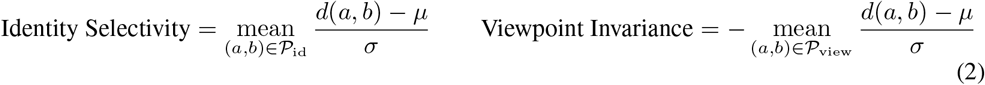

The viewpoint term is negated so that higher is better for both. Both indices are averaged over all 3 ^*N*^ three-identity subsamples.

### Estimating the viewpoint and identity subspaces

To estimate the viewpoint subspace, we average the embeddings over identities to obtain one mean per viewpoint. We center these means to remove the viewpoint-independent grand mean, and take an orthonormal basis *B* from the singular value decomposition of the centered means. The rank is capped at 2, as three centered means span at most two dimensions. The identity subspace is constructed similarly; we average the embeddings over viewpoints, center, and take the span. Both subspaces are fit onto three identities and evaluated on held-out identities.

### Two-dimensional embeddings

For the models in **Figure 2a, b** we visualize the 18 face-bank embeddings (6 identities × 3 viewpoints) in two dimensions. We z-score each feature across the 18 images so that a few high-variance identity dimensions do not dominate the geometry, L2-normalize each embedding, and take the matrix of pairwise cosine distances. Coordinates are the two-dimensional metric MDS on these distances; scale and orientation are arbitrary. The subspace-removed panels apply the same pipeline after projecting out the relevant subspace.

### Predicting recognition from the two indices

We fit an ordinary least-squares regression predicting each model’s CFMT score (number correct of 72) from its viewpoint-invariance and identity-selectivity indices, cfmt = *b*_0_ + *b*_1_ view + *b*_2_ id, across the 53 models. We report the model *R*^2^ and its together with each index’s individual Spearman correlation with CFMT score and the correlation between the two indices. **Figure 2c** plots the in-sample fitted values against the true scores.

### Removing a subspace

Removal is an orthogonal projection onto the complement of the subspace: each embedding *f* becomes *f* − (*f B*^*⊤*^)*B*, where the rows of *B* are the orthonormal basis. Removing the viewpoint subspace this way increases identity separation across models **Figure 2d**; the effect on the example embeddings is shown in **Figure 2a,b** (bottom).

### Ablating units

For each embedding dimension we compute the variance of its identity means (identity variance) and of its viewpoint means (viewpoint variance), on the fitting identities only, and rank units in descending order of (1) viewpoint variance (*viewpoint-dependent*), (2) identity variance (*identity-selective*), and (3) their ratio, identity over viewpoint (*viewpoint-invariance*).

The last two differ, since a unit can rank high on identity variance and low on the ratio. We ablate the top *k* units under each criterion by replacing each with its mean across the scored images, removing its information while leaving the scale of the embedding intact. We use *k* = 200 in **Figure 2f**, and show the sweep *k* from 1 to 200 **Figure 2e**. To check stability, we further measure the overlap of the two top-20 sets across disjoint halves of the identities, against the 20*/D* expected by chance.

### Technical Constraints

All embeddings and behavioral evaluation runs were performed on a single NVIDIA GPU (NVIDIA RTX A6000) in float32, and an AMD Ryzen Threadripper PRO 5975WX 32-core CPU. The full set of experiments completes in under 1 hour; no model training was required.

